# Corpus Callosum Dysgenesis impairs metacognition: evidence from multi-modality and multi-cohort replications

**DOI:** 10.64898/2026.03.02.709173

**Authors:** J.M. Barnby, R. Dean, H. Burgess, P. Dayan, L. J. Richards

## Abstract

The corpus callosum is the largest commissure in the mammalian brain and plays a major role in supporting cognitive processes required for adapting to complex environments. Individuals born with Corpus Callosum Dysgenesis (CCD), characterized by malformations of the corpus callosum, commonly exhibit deficits in social navigation, abstract problem-solving, decision-making, and self-awareness. Metacognition is a key cognitive process that supports these functions; however, it has yet to be tested comprehensively in individuals with CCD. Over three experiments, and three CCD cohorts, we tested the impact of this neurodevelopmental disorder on perceptual accuracy, confidence judgements, and metacognitive efficiency using two variants of a Random Dot Kinematogram task within lab, online, and VR conditions. We found that individuals with CCD typically displayed normal perceptual accuracy but failed to adjust their confidence judgements in line with task difficulty. Computational modelling revealed that this difference was explained by lower metacognitive efficiency driven by consistently lower metacognitive sensitivity. Together, these results provide evidence that the corpus callosum plays a crucial role in supporting metacognition.

## Introduction

Metacognition concerns the assessments individuals make about their own cognition (Fleming, 2024). Metacognition plays a critical role in adaptive functioning: it guides the gathering of information (Schulz et al., 2020), disclosure of personal information (Bang et al., 2020), assessment of personal knowledge during social interaction (Bahrami et al., 2012) and aids the regulation of physiological stress responses (Yoris et al., 2015). These processes depend on coordinated neural activity across several brain regions (Morales et al., 2018), especially during high cognitive demand (Fleming et al., 2024).

Metacognition is also implicated in psychiatric disorders (Seow et al., 2021), functional neurological disorders (Sadnicka et al., 2025), and Alzheimer’s disease (Bertrand et al., 2016). These disorders provide compelling examples of failures in neural circuits that support domain-specific and general metacognition (Morales et al., 2018; Vaccaro et al., 2018). One such population is those with corpus callosum dysgenesis (CCD; Paul et al., 2007; Edwards et al., 2014; Brown & Paul, 2019). CCD represents a continuum of neurodevelopmental disorders that stem from a congenital malformation of the corpus callosum. CCD includes a complete absence of callosal axons crossing the midline (agenesis), morphological changes in the anterior-posterior midline length (partial agenesis, leaving behind a callosal remnant of varying size), or thickness of the corpus callosum (hypoplasia and hyperplasia) (Paul et al., 2007, Glass et al., 2008).

Given the essential role of metacognitive monitoring to social and strategic inference (Frith, 2012), the partial overlap of neural regions encoding both (Vaccaro et al., 2018), and the centrality of impaired social inference in the CCD phenotype (Maxfield, et al., 2019; Maxfield et al., 2021), it begs the question as to whether the corpus callosum is vital to support metacognitive capability. CCD individuals commonly exhibit difficulties in neuropsychological assessments related to general metacognition, involving reflecting on, regulating, and guiding information processing across modalities. This includes learning, memory, and processing speed (Brown & Paul, 2019), abstract non-verbal reasoning (Hearne et al., 2019), and social deception (Barnby et al., 2022). Importantly, those with CCD have been consistently noted by clinicians and family members to underestimate their errors within reasoning and problem-solving tasks (Miller et al., 2024; Mangum et al., 2021).

We hypothesize that metacognition, a sophisticated higher-order cognitive process, will be vulnerable to the absence or malformation of the corpus callosum. Despite suggestive evidence, no systematic assessment has been conducted to investigate metacognition in CCD. Metacognition has also yet to be explored in the context of related anatomical changes such as callosotomy (cf: de Haan et al., 2020). In healthy volunteers, metacognitive capacity has been linked with a distribution network of regions, including the precuneus and ventromedial prefrontal cortex (Morales et al., 2018; Vaccaro et al. 2018; Zheng et al., 2021). Evidence is divided as to whether the corpus callosum may be vital in this coordination (Fleming et al., 2010). Calls have been made for further enquiry into whether the parallel functional coupling of neural regions, such as that afforded by the corpus callosum, is necessary for metacognition (Morales et al., 2018).

We conducted three experiments using variations of a well-validated Random Dot Kinematogram (RDK; Bang et al., 2018) task in three different cohorts of individuals with CCD. The RDK task is widely used in perceptual decision-making research as a canonical forced-choice paradigm. When coupled with confidence ratings, the task can be used to evaluate how accurately individuals calibrate their confidence to the accuracy of their choices. Computational frameworks have proposed that the relationship between metacognitive sensitivity (measured as a quantity known as meta-d’) and perceptual performance (the discriminability d’) can be cast as the efficiency by which confidence scores are deployed, taking account of true performance (meta-d’/d’; Maniscalco & Lau, 2012).

We administered the RDK to three cohorts of participants online, in the lab, or via a virtual reality headset; this latter modality allowed us to examine how the lateralization of visual input influences metacognition. We anticipated that individuals with CCD would exhibit an impaired ability to adjust their confidence ratings in response to changes in task difficulty, consistent with decreased metacognitive efficiency. Our results show evidence of robust and repeatable deficits in metacognition in individuals with CCD.

## Methods

### Participants

The University of Queensland Human Research Ethics Committee and Washington University St. Louis Institutional Review Board approved all procedures. All participants involved in the study provided informed consent to all research activities in this work. All CCD participants provided a neuroradiological scan or report to the research team as evidence of their diagnosis, unless otherwise specified. Participants were not included in the study if they had a diagnosis of a major neurological disorder, psychiatric illness, or epilepsy. Neurotypical controls were tested online (Experiment 1) or in the lab (Experiment 2 and 3). We provide further details of each sample below and report their demographics in Table 1.

**Table 1.** Group demographics across experiments.

|  | Experiment 1 (Online) |  | Experiment 2 (Lab) |  | Experiment 3 (VR) |  |
| --- | --- | --- | --- | --- | --- | --- |
|  | CCD | NT | CCD | NT | CCD | NT |
| N | 10 | 85 | 11 | 7 | 10 | 13 |
| Age<br>(mean[sd]) | 35.40<br>[15.32] | 28.20<br>[7.62] | 44.00<br>[19.59] | 42.71<br>[16.84] | 26.70<br>[7.73] | 29.69<br>[10.31] |
| Sex<br>(m:f:nb) | 4:6:0 | 34:50:1 | 3:8:0 | 2:5:0 | 8:2:0 | 6:7:0 |

### Random Dot Kinematogram (RDK)

Two variants of the RDK task were used. The first (experiments 1 and 2) used a conventional version of a Random Dot Kinematogram task inspired by previous work (Rausch & Zehetleitner, 2014, Bang et al., 2018), which evaluates baseline visual perception and confidence estimates. During implementation of this version, feedback was sought from a CCD participant via a short pilot study. In response, a set of simplifications were made to the reference presentation and confidence input mechanisms (see Supplementary Materials for modification details).

The second variant (experiment 3) further modified the stimulus so that it could be restricted to a single eye and/or visual hemifield to better understand the effects of perceptual laterality on confidence judgements in these individuals (see Supplementary Materials for modification details). This modified version of the RDK task was conducted within a virtual reality environment.

To summarize the timelines of both tasks (see below for details on each individual setup), the participants were first taken through a tutorial, with instructions provided in writing and verbally by an experimenter, followed by 20 practice trials to familiarize themselves with the task and the mechanism for providing their responses. The participants then undertook a set of calibration trials, which we used to normalize the difficulty of the task across participants using a ‘two-up-one-down’ staircasing procedure, before moving on to the main trials. Participants completed 120 calibration trials, in the first of which 20% of the dots moved coherently. Two correct judgments in succession decreased the coherence percentage by 1%, while a single incorrect judgment increased the coherence percentage by the same value; the coherence percentage was constrained to lie between 12% and 50%. From these calibration trials a fitted coherence parameter, *K*_med_, was derived for each participant to create a set of taking the median of the coherence values from the final 30 trials of the calibration phase. This value was doubled (×2) and halved (×0.5) to generate the pair of high and low moving dot coherence percentages the participant would experience during the main trials. For the modified version of the RDK task in experiment 3, a different *K*_med_ was calculated for each of the three different forms of stimulus presentation (i.e., binocular, monocular, and lateralized; see below for further details). After calibration, each participant moved to the main phase of the task, in which they performed a further 200 trials, with the dot coherence alternating randomly between the calibrated high and low moving dot coherence percentages.

For each task version, participants were presented with instructions prior to the initial calibration trials describing the typical composition of a trial: presentation of dot motion within an aperture followed by a decision based on the motion they observed, including a confidence judgement based on what best represented their conclusion. These instructions included a preview of the dot motion and a screen for practicing making input selections. We emphasized that participants should identify the motion of the coherent dots rather than the overall dot motion. Participants were then informed of how many calibration trials they would complete before starting the trials. After completing the calibration trials, participants were shown instructions notifying them of their progress and that they were proceeding to the main trials of the experiment. At the conclusion of the experiment, participants were notified of the experiment’s completion.

Participants were excluded from the metacognitive component of the study if the adjusted dot coherence percentage would have exceeded 50% at any point during the calibration trials, if they made more than 20 mistakes during the main trials (i.e., more than 10%), or if there was no variance in their self-reported confidence.

#### Experiment 1

The purpose of this experiment was to establish initial evidence for differences between the metacognitive efficiency, perceptual accuracy, and decision confidence of individuals with CCD and neurotypical (NT) controls. An online version of the RDK task, written in JavaScript (see Code & Data for a link to all open materials), was delivered to participants via Gorilla.sc. 11 participants with CCD (see supplement for exact callosal malformation; one participant was unable to provide a radiological scan and self-reported their diagnosis to the team) were invited to undertake this task as members of a US-based non-profit organization for individuals with a diagnosis of CCD and their families; one participant was excluded due to non-convergence of calibration trials. All NT participants for this experiment were recruited via Prolific.ac. In addition to the exclusion criteria for CCD individuals, online participants were screened out if they were under the age of 18 or over the age of 65, if they had a lower than 90% approval rating on Prolific, or if they were currently receiving any prescribed medication. A total of 86 NT participants were invited to participate; one was excluded due to a failed calibration procedure.

The task itself was a modified version of the Random Dot Kinematogram task that blends perceptual judgements with confidence estimates, such as that used within Rausch & Zehetleitner, 2014 (see Figure 1). On each trial, participants were presented with a circular aperture, which was 8.0 degrees in diameter, within which a field of moving dots was visible; each dot (0.12 degrees in diameter) was visible for 16.67 ms on a 60 Hz display before being replotted. The field consisted of two independent sets of dots, 1) those that moved coherently and 2) those that moved randomly; the percentage of coherently moving dots was determined by the motion coherence. The coherently moving dots were displaced in the target direction, either left or right (parallel to the horizontal meridian), at a rate of 2 degrees s^-1^, whereas each of the randomly moving dots was displaced at 2 degrees s^-1^ in a random direction (which was fixed at the time of their creation). During dot motion, the average density of visible dots was 12.5 dot degrees^-2^ s^-1^. To preserve the visible dot density, dots outside the aperture were hidden, and upon reaching a display canvas boundary their position would be reset to the opposite boundary of the same axis. To assist the participants in maintaining fixation, a circular region (0.6 degrees in diameter) at the center of the aperture was kept free of dots with a cross (0.4 degrees in diameter) at its center. The moving dots were visible to the participants for a total of 1.5 s; after the dots were hidden, the participant was required to give either a ‘left’ or ‘right’ response via keyboard, using their respective left or right index finger, before the next trial commenced. In keeping with prior work (Bang et al., 2018), every five trials, participants were required to rate their confidence as to whether they thought they had correctly judged the target direction of the coherently moving dots in the most recent trial. Participants rated their confidence via keyboard, using their left index finger to decrease their rating and their right index finger to increase it, via a visual analogue scale that ranged from 50% to 100% with 10% increments.

**Figure 1.**
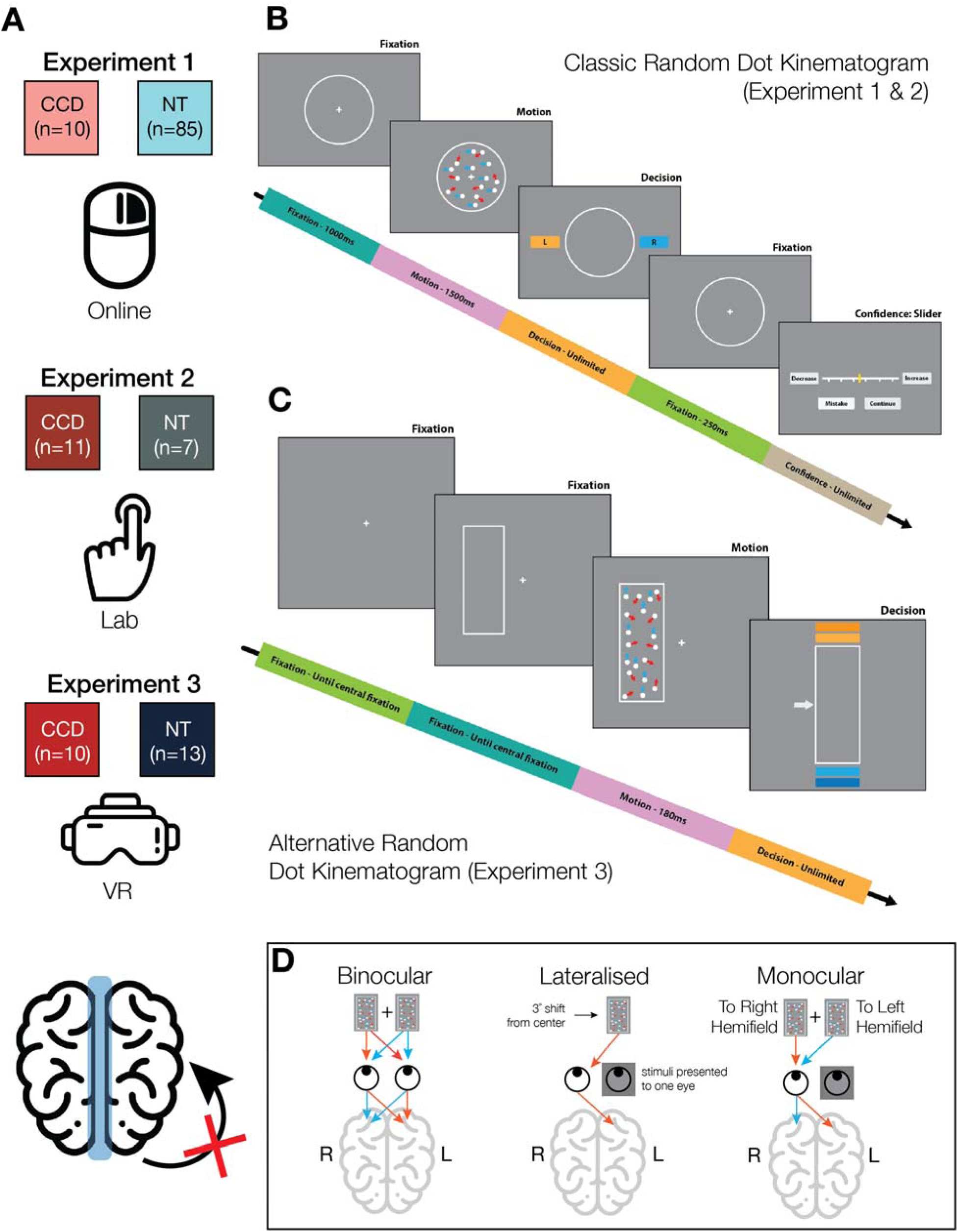
Experiment Summary. We hypothesized that malformations or absence of the corpus callosum would impair metacognition. (A) The study consisted of three experiments, testing three different participant samples that included individuals with a diagnosis of corpus callosum dysgenesis (CCD) and neurotypical controls (NT). Experiment 1 was performed online, while Experiment 2 was conducted in the lab with matched controls, and Experiment 3 within a virtual reality (VR) environment. (B) Experiment 1 and 2 used a classic Random Dot Kinematogram (RDK) task that has been previously described (Bang et al., 2018). This task was implemented using JavaScript (Burgess et al., 2023) and hosted on Gorilla.sc. (C) The RDK task used in the VR environment was modified to require participants to simultaneously record the perceived dot direction and their confidence in this decision (see Methods for further details). (D) The Modified RDK from Experiment 3 in VR allowed for Binocular (both hemifields to both eyes), Lateralized (isolated hemifield to one eye), and Monocular (both hemifields to one eye) manipulations.

Analysis was conducted using R (v4.3.3). All behavioral outcomes and confidence ratings were analyzed using random effects models with participant ID as a random variable. We extracted and reported t-values, effect sizes, and p-values from the outcomes of these models. We retained the average findings across groups for visualization. The behavioral data were processed to extract a latent measure of metacognitive efficiency (meta-d’/d’; Maniscalco & Lau, 2012). This involved binning the observed count of confidence attributions from 50-100 (in bins of 10) for each participant when perceptual decisions were correct and incorrect (see Figure S5). We then fitted the data using a model of metacognitive efficiency (HMeta-D; Fleming & Daw, 2017), which uses Bayesian Hierarchical fitting to estimate group and individual level parameters of metacognitive efficiency (herein, and within the model implementation, called the M-ratio, or *μ*_ratio_). We fitted the CCD and NT groups separately to extract group-level *μ*_ratio_, ratio, meta- *d*′ and *d*′ values. Of note, we jointly fitted both conditions together (*k*_med_ × 0.5 and *k*_med_ × 2.). We did this to uncover the overall difference in metacognitive efficiency between groups. We acknowledge that this can lead to inflated meta-*d′* estimates (Rahnev & Fleming, 2019) and report separate distributions of individual meta-*d′* and *μ*_ratio_ values for each condition for clarity (Figure S5). We were unable to do this for Experiment 3 due to the small number of trials per participant per condition. We report posterior predictive checks on confidence estimates and recovery analyses for *μ*_ratio_ values.

To assess the strength of evidence for the observed difference between groups we performed permutation testing of the group-level difference in metacognitive efficiency (Δ*μ*_ratio_), metacognitive sensitivity (Δmeta-*d′*) and perceptual sensitivity (Δ*d′*). This involved shuffling groups and refitting the models on each shuffled group to generate random group-level estimates. This was repeated 500 times to generate a null distribution representing a collection of values of Δ*μ*_ratio_ that may have been generated by chance. Cohen’ *U* and 95% Wilson confidence intervals (95%CI[*U*]) were then extracted. *U* is the strength of evidence that the observed Δ*μ*_ratio_*^obs^* is outside the permuted Δ*μ*_ratio_^*perm*^ values [*U* = Pr (Δ*μ*_ratio_^*obs*^ ≥ *μ*_ratio_^*perm*^) and does not assume normality or equal variance of distributions (Pastore & Calcagnì, 2019). *U* ≈ 0.50 means difference is superior to the null. 95%CI[*U*] were then calculated as follows: that a true observed difference is approximately the median of the null distribution values of 0.5 < *U* < 1 indicates the strength of evidence that the true observed difference is superior to the null. 95%CI[*U*], and were then calculated as follows:

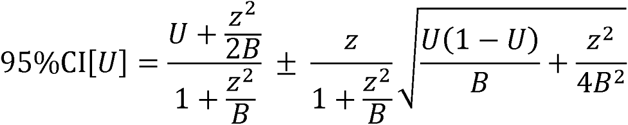

where B equals the number of permutations (in our case, 500) and *z* is 1.96 for a 95%CI. *U* and 95%CI[*U*] thus provide compact metrics to gauge the strength and uncertainty of evidence for each group difference across all experiments.

We report reaction time changes as a function of dot coherence and correctness within the supplementary materials.

#### Experiment 2

Here we sought to replicate the findings of Experiment 1 in the lab. We recruited 14 different individuals with CCD from an Australian non-profit organization for individuals with a diagnosis of CCD and their families and 7 neurotypical controls with comparable age and sex distributions (see Supplementary Materials). These participants undertook the RDK task described in **Experiment 1** whilst simultaneously undergoing functional magnetic resonance imaging (fMRI). All CCD participants provided radiological evidence of their diagnosis. The RDK stimulus (as described in Experiment 1) was projected onto a screen positioned outside the MRI spectrometer; participants viewed this screen using a pair of mirrored goggles. Participant responses were recorded using a four-button MR-safe response pad. The left thumb was used to submit ’left’ directional choices and decrease reported confidence ratings, while the right thumb was used to submit ’right’ directional choices and increase confidence ratings.

All behavioral and computational analysis was repeated identically to **Experiment 1**, and the same exclusion criteria were adhered to. 3 CCD participants were consequently excluded due to making more than 20 mistakes during the main trials and/or failing calibration procedures, leaving 11 (4 with complete and 6 with partial agenesis and 1 with callosal hypoplasia). The high degree of heterogeneity of CCD brain topologies significantly complicates the analysis of imaging data; we will therefore report this in a separate publication.

#### Experiment 3

The purpose of this experiment was to replicate our behavioral results in a new task environment that allowed for laterality manipulations of visual stimuli. 10 new individuals with CCD were recruited as members of a US-based CCD non-profit organization to undertake the VR version of the RDK task; 4 were unable to provide radiological reports and self-reported their diagnosis to the research team. 13 age and sex-matched neurotypical controls were recruited to provide a comparison. To maintain control over the visual delivery, participants undertook this task in virtual reality (VR) using a Meta Quest Pro headset. The Meta Quest Pro eye-tracking system was used to monitor the participants’ gaze during the task and actively adjust the position of the aperture to maintain fixation in real-time. The RDK stimulus was modified to suit the restricted visual environment of the VR headset by converting the apertures to vertically elongated rectangles (8.0 by 16.0 degrees) and adjusting the target directions of the coherently moving dots to either up or down, parallel to the vertical meridian. During dot motion, the average density of visible dots within the aperture was 16 dot degrees^-2^ s^-1^ (see Figure 1). We tested three different stimulus presentation conditions: binocular, when it was in the central visual field of both eyes; lateralized, when it was only shown in the nasal visual hemifield of a single eye; and monocular, when it was placed centrally, but only visible to a single eye (Figure 1D). During lateralized presentation trials, the aperture was offset so that the midpoint of its medial edge was 3.0° from the point of central fixation after accounting for the aperture width and inter-pupil distance (IPD).

Lateralizing the presentation of the moving dots was a natural extension of the previous two experiments; due to reduced interhemispheric connectivity, access to contralateral cortical resources for visual monitoring, processing, and evaluation was anticipated to be significantly curtailed in the CCD population. By restricting visual information to a single visual hemifield we would test the capacity of the CCD participants to use alternative means to recruit the necessary resources to maintain performance accuracy and metacognitive sensitivity. The monocular presentations, on the other hand, were used to test for the effect of a binocular advantage in individuals with CCD. The binocular integration of visual information has been hypothesized to be primarily attributable to the splenium of the corpus callosum (Mitchell and Blakemore,1970; Saint-Amour et al., 2004). In the absence of the splenium, binocular presentations of stimuli may not afford individuals with CCD a significant advantage in movement perception, which would be detectable via comparisons of performance between bilateral and monocular trials.

The mechanism through which participants made their responses was also modified. Instead of having separate reports for dot motion direction and confidence, the two were combined in a four-point Likert scale, consisting of the options ‘very confident of upwards motion’, ‘somewhat confident of upwards motion’, ‘somewhat confident of downwards motion’, and ‘very confident of downwards motion’. This change allowed for an assessment of confidence to be made in each trial and forced participants to give a non-neutral rating of confidence (albeit at the expense of granularity in confidence judgments). Participants used a Meta Quest Pro controller, held in their right hand, to record their responses. They manipulated the controller’s joystick with their right thumb to navigate between different response options and used their right index finger to hold down the controller trigger to submit their choices. For a response to be recorded, the controller trigger had to be held down for a full second.

All behavioral and computational analyses were conducted identically to **Experiment 1**. In contrast to the classic experimental task, only low or high confidence (1 or 2) could be reported for each dot direction (up or down) in the alternative version. These confidence ratings were scaled for visualization but were used as ordinal variables for modelling.

## Results

We sought to test the hypothesis that malformations or absence of the corpus callosum would impair metacognition (Figure 1). We predicted that this would be expressed by poor metacognitive sensitivity and metacognitive efficiency within Random Dot Kinematogram (RDK) tasks compared to neurotypical (NT) controls. Over three experiments we recruited groups of CCD and NT participants and tested them in online, lab-based, and VR-based environments (Figure 1A). **Experiments 1** and **2** involved a classic RDK task (Bang et al., 2018; Figure 1B), whereas **Experiment 3** involved an alternative RDK task (Figure 1C) designed to instill visual laterality processing biases depending on the hemifield presentation (Figure 1D).

### Participant demographics

After exclusion, the final participant counts for each task were: **Experiment 1**: unmodified RDK (online) - 10 CCD and 85 NT; **Experiment 2**: unmodified RDK (Lab) - 11 CCD and 7 NT; **Experiment 3**: modified RDK (VR) - 10 CCD and 13 NT (Table 1; Figure 1). For more information, see the corresponding description of the experiment in the Methods.

After exclusions, calibration within experiments was equivalent between groups (**Table S1**). There were between-experiment impacts of testing environment on calibration (**Table S1**); online experiments (Experiment 1) led to lower coherence values across the board.

### Experiment 1

We first observed that those with CCD are metacognitively less efficient than neurotypical controls in an online RDK task. We tested raw perceptual and confidence judgement differences between CCD and neurotypical participants who completed the unmodified RDK task online, assessing both behavioral accuracy and confidence, and metacognitive efficiency. Importantly, the staircase procedures were approximately equivalent between groups (see Supplementary Materials), reducing the likelihood that the following outcomes could be attributed to differences in individual difficulty.

The task showed good discriminability, such that higher moving dot coherence led to improved accuracy scores in both groups (CCD: t=8.33, es=0.77, p<0.001; NT: t=15.47, es=0.48, p<0.001). There was a group by coherence interaction, such that NT participants were more accurate than CCD participants in low but not high coherence trials (t=3.29, es=0.31, p=0.001; see Figure 2A).

**Figure 2.**
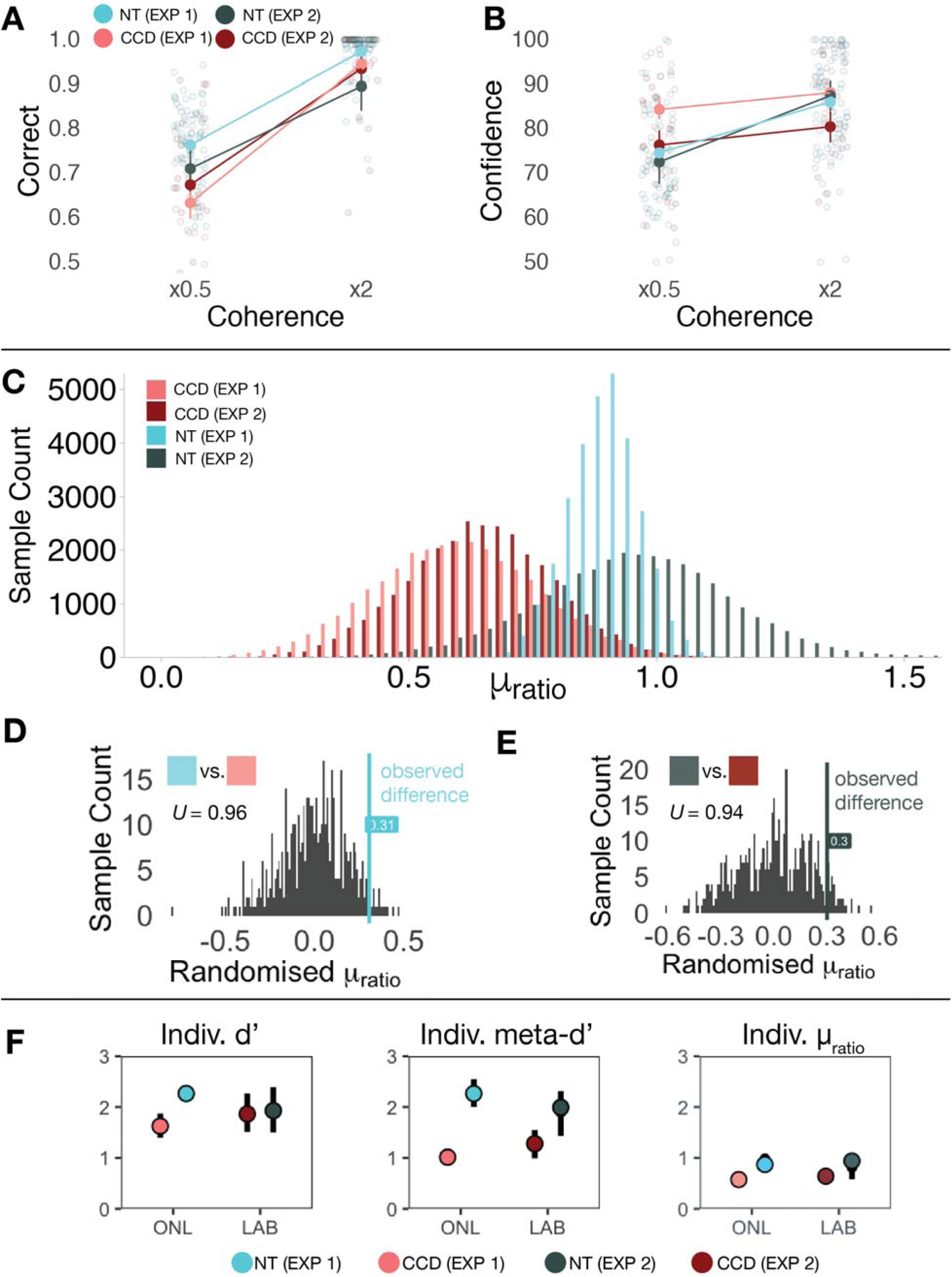
Behavioral and Computational Results for Experiment 1 and 2. (A) Average perceptual accuracy for both CCD and NT groups for Experiments 1 and 2; both groups showed improvements in accuracy in line with stronger coherence (*k*_med_ × 0.5 and *k*_med_ × 2). (B) Average confidence ratings for both CCD and NT groups for Experiments 1 and 2; CCD groups showed consistent failures to calibrate confidence scores in line with changing coherence. (C) Distribution of group-level *μ*_ratio_ for both CCD and NT groups within Experiments 1 and 2; both CCD groups showed lower group-level *μ*_ratio_ values than NT participants. (D). Permutation analysis tested whether the observed difference in group-level metacognitive efficiency parameters (Δ*μ*_ratio_) were whether the observed difference in group-level metacognitive efficiency parameters (*μ*_ratio_) were within the null distribution in Experiment 2; this showed good evidence of Δ*μ*_ratio_ being outside the null (*U*=0.94; 95%CI[*U*]: 0.92, 0.96). (F) 95% confidence intervals and hierarchical group mean (colored dot) representing individual values of *d′*, meta-*d′*, and *μ*_ratio_ for CCD and NT participants across Experiment 1 and 2.

Both CCD and NT participants increased their mean confidence in line with increasing coherence (CCD: t=4.20, es=0.35, p<0.001; NT: t=27.51, es=0.69, p<0.001). There was a group by coherence interaction, such that NT participants had lower confidence in lower coherence conditions versus CCD participants (t=6.23, es=0.47, p<0.001; Figure 2B).

To assess the metacognitive capability of the participants, we calculated metacognitive efficiency (*μ*_ratio_) using a Bayesian hierarchical procedure (Fleming & Daw, 2017). This allowed us to evaluate whether participants scaled their confidence (meta-d’) in line with their reported perceptual accuracy (d’). A permutation analysis (500 repetitions) was conducted to assess whether the true difference between group-level _ratio_ mean values lay outside of the null distribution.

Computational modelling revealed that those with CCD were far less efficient in their calibration of confidence relative to their perceptual accuracy (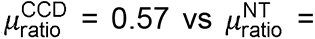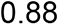; Figure 2C). There was strong evidence that the difference in distributions was outside the null (*μ*_ratio_ = 0.31, *U*=0.96, 95%CI[*U*]: 0.94, 0.98; Figure 2D). Group differences in efficiency were driven primarily by large reductions in metacognitive sensitivity (Δmeta-*d*′=1.29, *U*=0.99, 95%CI[*U*]: 0.98, 0.99) in CCD compared to NT participants (Figure 2F). Smaller differences in perceptual sensitivity were also found (Δ*d*′=0.64, *U*_3_=0.99, 95%CI[*U*]: 0.98, 0.99).

The model was able to successfully produce data that strongly correlated with observed values for each confidence bin in both left and right dot directions (pearson r range = 0.557-0.995, ps < 0.01); *μ*_ratio_ values were strongly correlated for both CCD (r=0.75, p=0.02) and NT participants (r=0.91, p<0.01).

Within our online neurotypical cohort, we also administered the ICAR progressive matrices to measure abstract non-verbal reasoning (Condon & Revelle, 2014). The average ICAR score within our neurotypical sample was 6.79±2.28 (typical general population score [n=79801] is 5.23±0.47; Condon & Revelle, 2014). There was no correlation between the ICAR and 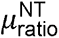 (r =0.050, p=0.65), *d*′(r=-0.064, p=0.56) or meta-*d*′ (r=0.01, p=0.96).

### Experiment 2

The primary findings of Experiment 1 replicated when the RDK task was administered to participants in the lab with matched controls. As with Experiment 1, staircase procedures led to equivalent difficulty between groups (see Supplementary Materials). As found in Experiment 1, the task showed good discriminability, such that higher dot coherence led to improved accuracy scores in both groups (CCD: t=7.41, es=0.68, p<0.001; NT: t=4.13, es=0.47, p<0.001; Figure 2A; Figure S3) with the exception that there was no interaction of group by coherence. Furthermore, both CCD and NT participants increased their mean confidence in line with increasing coherence (CCD: t=3.95, es=0.28, p<0.001; NT: t=10.30, es=0.90, p<0.001; Figure 2B). Once again, there was an interaction between group and coherence on confidence scores in the same direction as Experiment 1 (t=6.18, es=0.70, p<0.001; see Figure 2B).

Computational modelling revealed that those with CCD were less efficient in their calibration of confidence relative to their perceptual accuracy (^CCD^ = 0.63 vs ^NT^ = 0.94; Figure 2C), although with a broader group distribution than Experiment 1. There 95%CI[*U*]: 0.92, 0.96; Figure 2E). Differences in efficiency were primarily driven by reductions in metacognitive sensitivity in CCD participants, although evidence for this reduction was weaker (meta- =0.71, =0.89, 95%CI[*U*]: 0.86, 0.91) than in experiment 1. *d′* was comparable between groups with no evidence for difference from the null (Δ*d*′=0.07, *U*=0.54, 95%CI[*U*]: 0.49, 0.58; Figure 2F).

The model was able to successfully produce data that strongly correlated with observed values for each confidence bin in both left and right dot directions (pearson r range = 0.802-0.996, ps < 0.01); _ratio_ values were correlated for both CCD (r=0.59, p=0.06) and NT participants (r=0.86, p=0.02).

Within our lab-based CCD cohort, we also had data on 9/11 participants who had completed the ICAR progressive matrices to measure abstract non-verbal reasoning (Condon & Revelle, 2014). The average ICAR score in this subsample was 4.78±2.73 which is not significantly different from the general population average previously reported in Condon & Revelle, 2014 (one-sample t=-0.49, p=0.63). There was no correlation between the ICAR and 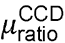 (r =0.52, p=0.15), *d*′(r=-0.39, p=0.30) or meta- *d′*(r=0.45, p=0.22).

### Experiment 3

The behavioral and computational results were replicated for a third time when participant performance was assessed on the binocular trials of the VR modified version of the RDK task. During this version of the RDK task, the stimulus alternated between three presentation modes: binocular, monocular, and lateralized (restricted to a single nasal visual hemifield). **Table 2** summarizes the group-average accuracy rates, the percentage of high-confidence reports, and average reaction times for these different forms of stimulus presentation. As with **Experiments 1** and **2**, staircase procedures led to equivalent difficulty between groups but with some important differences between conditions (see Supplementary Materials).

**Table 2.** Mean group accuracy and reaction times given stimulus presentation in the modified RDK task (VR environment).

|  | CCD |  |  |  | NT |  |  |  |
| --- | --- | --- | --- | --- | --- | --- | --- | --- |
|  | Low Coherence |  | High Coherence |  | Low Coherence |  | High Coherence |  |
| Presentation | Accuracy (%[sd]) | RT (s[sd]) | Accuracy (%[sd]) | RT (s[sd]) | Accuracy (%[sd]) | RT (s[sd]) | Accuracy (%[sd]) | RT (s[sd]) |
| Binocular | 61.2<br>[16.9] | 2.79<br>[1.53] | 77.0<br>[19.3] | 2.68<br>[1.27] | 66.1<br>[13.9] | 2.30<br>[0.56] | 84.9<br>[14.7] | 2.07<br>[0.40] |
| Monocular | 53.0<br>[14.1] | 2.73<br>[1.20] | 70.0<br>[15.8] | 2.59<br>[0.80] | 67.0<br>[10.2] | 2.08<br>[0.33] | 83.4<br>[19.5] | 2.14<br>[0.47] |
| Lateralized | 55.0<br>[14.7] | 3.11<br>[1.93] | 63.1<br>[12.2] | 2.76<br>[1.35] | 58.5<br>[13.5] | 2.42<br>[0.81] | 76.3<br>[17.4] | 2.16<br>[0.47] |

#### Behavioral Analysis

The VR-based task showed good discriminability in binocular trials, such that higher dot coherence led to improved accuracy scores in both groups (CCD: t=2.68, es=0.24, p=0.008; NT: t=3.91, es=0.30, p<0.001; Figure 3A). Analysis of lateralized trials showed worse performance of CCD individuals, but equivalent performance of NTs (CCD: t=1.90, p=0.058; NT: t=4.89, es=0.27, p<0.001). Analysis of monocular trials showed accuracy improved when coherence increased in both CCD and NT individuals (CCD: t=3.88, es=0.24, p<0.001; NT: t=4.91, es=0.27, p<0.001). There was no group interaction with coherence within any presentation type in Experiment 3. Analysis of all data together, irrespective of presentation, using random-effect models type found main effects for coherence (t=5.12, es = 0.21, p<0.001) and group (t=2.30, es=0.21, p=0.03) on accuracy.

**Figure 3.**
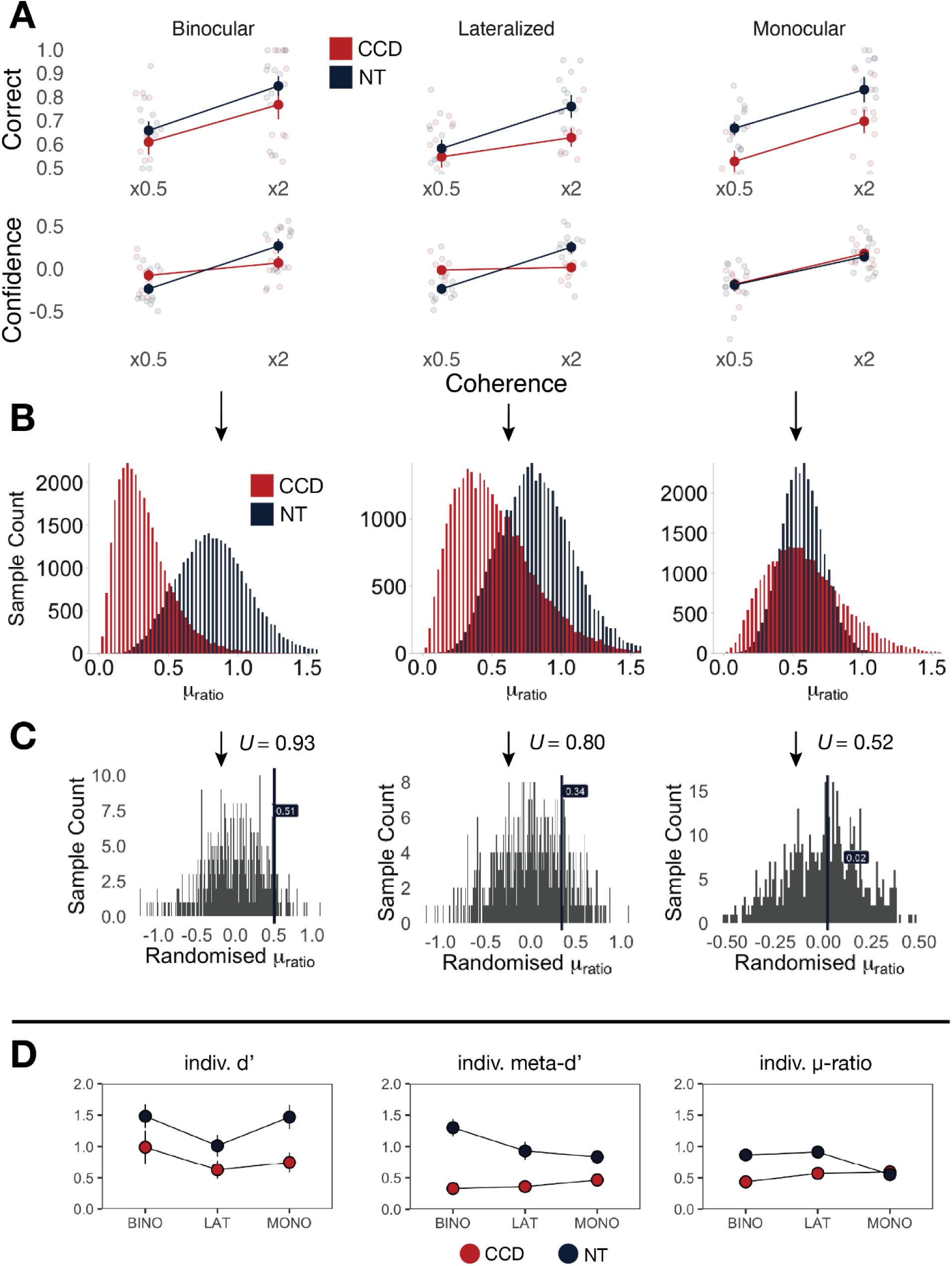
Behavioral & Computational Results from Experiment 3. (A) Average perceptual (top) and confidence (bottom) ratings for both CCD and NT groups within binocular, lateralized, and monocular trials recorded during Experiment 3. (B) Distribution of group-level *μ*_ratio_ for both CCD and NT groups within binocular, lateralized, and monocular trials recorded within Experiment 3. (C). Permutation analyses (500 repetitions) indicate that binocular trials showed a low probability of observed differences group-wise in efficiency parameters (Δ*μ*_ratio_) being within the null distribution (p=0.048), whereas lateralized and monocular trials did not (p=0.312; p=0.480). (D) 95% confidence intervals (black bar) and hierarchical group mean (colored dot) representing individual values of *d′*, meta-*d′*, and *μ*_ratio_ for CCD and NT participants across binocular (BINO), monocular (MONO), and lateralized (LAT) experimental conditions.

In binocular trials, those with CCD did not change their mean confidence in line with higher dot coherence, whereas NT participants did (CCD: t=1.07, p=0.29; NT: t=4.25, es=0.36, p<0.001; Figure 3A). There was an interaction between group and coherence (t=2.01, es=0.26, p=0.045). This was also true for lateralized trials (CCD: t=0.30, p=0.76; NT: t=5.79, es=0.35, p<0.001; group × coherence: t=3.60, es=0.33, p<0.001). Interestingly, both CCD and NT participants adjusted their confidence in line with coherence during monocular trials (CCD: t=3.89, es=0.25, p<0.001; NT: t=4.09, es=0.23, p<0.001). There was no group-by-coherence interaction for monocular trials. For raw paired differences between conditions, see Figure S4. Analysis of all data together, irrespective of presentation, using random-effect models type found main effects for coherence (t=3.21, es = 0.13, p=0.001) and a group by coherence interaction (t=2.93, es=0.16, p=0.003) on confidence, such that neurotypical participants reduced their confidence more on lower coherence trial and increased their confidence more on higher coherence trials (see Figure 3).

Reaction times were faster in NT participants versus CCD participants (t=-4.93, es=-0.29, p<0.001) and were faster in binocular versus lateralized trials (t=-2.58, es=0.14, p=0.010). There was no interaction between group and presentation type. We report a detailed analysis of reaction times by dot coherence and correctness in the **Supplementary Materials.**

#### Computational Modelling

Binocular trials were presented to be functionally identical to stimulus presentations within **Experiments 1** and **2**.

Computational modelling revealed that those with CCD were far less efficient in their 0.28 vs in their calibration of confidence relative to their perceptual accuracy in binocular trials 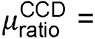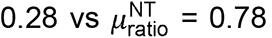 Figure 3B). Permutation analysis showed good evidence that this observed difference was outside the null (Δ*μ*_ratio_ =0.50, *U*=0.93, 95%CI[*U*]: 0.90, 0.95; Figure 3C). The difference in efficiency was driven by large differences in metacognitive sensitivity (Δmeta-*d*′=1.10, *U*=0.96, 95%CI[*U*]: 0.93, 0.97) between groups (Figure 3D). There was also good evidence for a smaller difference in perceptual sensitivity (Δ*d*′=0.49, *U*=0.93, 95%CI[*U*]: 0.91, 0.95)

Compared to binocular trials there was a smaller difference in efficiency between groups for lateralized and monocular trials. This was due to numerically equivalent or worsening efficiency in NT participants (Lateralized: 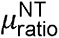 = 0.79; Monocular 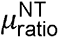 = 0.54), and numerically improved efficiency in CCD participants (Lateralized: 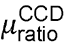 = 0.44; Monocular; 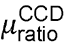 = 0.51). There was weak evidence for between-group differences existing outside of the null in both Lateralized and Monocular conditions (Lateralized: Δ*μ*_ratio_=0.34, *U*=0.80, 95%CI[*U*]: 0.76, 0.83; Monocular: Δ*μ*_ratio_=0.02, *U*_3_=0.52, 95%CI[*U*]: 0.48, 0.56).

The changes in efficiency between groups in lateralized and monocular trials were driven by two factors: perceptual sensitivity was higher in NT compared to CCD participants in lateralized trials (Δ*d*′=0.47, *U*=0.95, 95%CI[*U*]: 0.93, 0.97), and this difference between groups increased in monocular trials (Δ*d*′=0.82, *U*=0.99, 95%CI[*U*]: 0.98, 0.99); Metacognitive sensitivity was higher in NT compared to CCD participants in in monocular trials (Δmeta-*d*′=0.44, *U*=0.95, 95%CI[*U*]: 0.92, 0.96). Perceptual and metacognitive sensitivity parameters within participants and between conditions were all correlated (=0.51-0.75, ps<0.05; see Supplementary Materials).

The model was able to successfully produce data that strongly correlated with observed values for each confidence bin in both left and right dot directions (pearson r range = 0.885-0.978, ps < 0.01); *μ*_ratio_ values were correlated for both CCD (r=0.79, p<0.01) and NT participants (r=0.57, p=0.045).

In sum, the findings of **Experiments 1** and **2** were replicated in binocular VR trials, demonstrating that those with CCD have poorer metacognitive efficiency compared to NT participants, primarily driven by worse metacognitive sensitivity. The results of our visual manipulations suggest that the seemingly higher metacognitive efficiency of CCD individuals in lateralized and monocular conditions is attributable to worsening perceptual sensitivity compared to binocular trials.

## Discussion

We tested the importance of cortical interhemispheric connectivity in metacognitive processing by examining individuals with a diagnosis of CCD. We assessed perceptual accuracy and decision confidence in both CCD and NT participants using two variants of the RDK task across three experiments.

Perceptual accuracy was approximately equivalent between the two groups. Sensitivity analysis showed robust effects irrespective of staircasing, task variation, and task delivery. This broadly conforms with past work, which has shown that in simple perceptual tasks, those with CCD can perform on average in line with neurotypical controls (Brown & Paul, 2019). Indeed, the CCD participants performed the tasks at similar levels of difficulty to their NT counterparts.

Nevertheless, those with CCD did not benefit from higher coherence when calibrating confidence ratings; when the task became easier, confidence did not increase, unlike for the NT controls. Previous work has shown that, in neuropsychological testing environments, those with CCD are less able to notice their errors in problem-solving and reasoning tasks (Mangum et al., 2021), errors which are otherwise evident to family members (Miller et al., 2024). This ‘flattening’ of absolute confidence values was higher on average in online-based CCD participants but still showed a relative insensitivity to changing task coherence.

Behavioral findings were borne out in computational quantification of metacognitive efficiency (Maniscalco & Lau, 2012). Using a well-established model of implementation of metacognitive efficiency (Fleming, 2017) that we fit to perceptual and confidence data, we determined a consistent reduction in efficiency. This was driven by lower metacognitive sensitivity, in those with CCD versus NTs, irrespective of task variation and delivery.

These results prompt two questions: what is the typical role of the corpus callosum in supporting neurotypical metacognition? And what does this tell us about the neural ‘substrate’ of metacognition?

Nonsystematic causes of metacognitive inefficiency have been suggested to be due to inherent processing or resource limitations (Shekhar & Rahnev, 2021; Maniscalco & Lau, 2015; Maniscalco et al., 2017). Previous evidence suggests that domain-specific and domain-general metacognition involve distinct processes, the former relying on anterior prefrontal encoding, with the latter involving a distributed network of frontal and posterior midline regions (Fleming et al., 2010; Morales et al., 2018; Vaccaro et al., 2018). The role and importance of the corpus callosum in facilitating the coordination of these regions to support metacognition has been mixed (Fleming et al., 2010; Zheng et al., 2021).

We hypothesize a proximal explanation: domain-general metacognition partly depends on the cognitive efficiency afforded by the corpus callosum, and thus the corpus callosum represents a crucial bottleneck of cognitive resources that are essential for metacognitive function. This hypothesis is further supported by reaction time data (those with CCD are slower to respond; see Supplementary Materials), and the results of our laterality manipulations of the RDK task in VR. However, demonstrating this requires separation of perceptual and metacognitive sensitivities.

We examine our findings considering systematic and nonsystematic effects that may cause metacognitive inefficiency (Shekhar & Rahnev, 2021). When both cortices can be engaged through binocular presentations across all experiments, perceptual sensitivity is at its highest, with both groups demonstrating sensitivity to dot coherence changes. The decrease in perceptual sensitivity during the lateralized condition suggests that both CCD and NT individuals rely on input to both visual hemifields to accurately gauge the motion of moving dots. Likewise, the results from the monocular presentations indicate that both groups benefit from binocular input. This implies that perceptual sensitivity was more dependent on the total visual information available. Thus, under binocular conditions, differences in metacognitive efficiency between groups are less likely to be due to failures in sensory input.

Metacognitive sensitivity, on the other hand, is far more brittle. Indeed, across all conditions, metacognitive sensitivity is overall lower in CCD compared to NT participants. This strengthens the argument that metacognitive efficiency differences in binocular conditions seen across experiments are due to nonsystematic computational failure, which we hypothesize is due to bottlenecks in information integration across the brain midline. Our introduction of multiple task variants weakens systematic explanations, which could be due to the way confidence is measured or recorded, although it does not overcome the validity of lab-based measurement versus real-world deployment of confidence (Rahnev, 2020). The decrease in metacognitive sensitivity in NT participants during monocular trials compared to binocular trials, despite higher perceptual sensitivity, suggests that cognitive monitoring cannot be stabilized in informationally incomplete environments, and is thus a failure of computation, not input, even with a fully formed corpus callosum. Systematic failures in monocular and lateralized trials cannot be fully ruled out, as they were only included in one experiment.

A potential confounder of our hypothesis is that a deficit in broader cognitive functions may bias the metacognitive outcomes reported here. However, we consider this to be an unlikely explanation for our findings. Wright & Booth (2023) noted that of the studies that report on the general intelligence of those with CCD, more than half reported that their cohorts scored within the normal range (7/13). Our own prior work (Barnby et al., 2022; Hearne et al., 2019) also demonstrates that those with CCD do not struggle with abstract non-verbal reasoning tasks compared with neurotypical participants, except when items are very complex. We additionally found no relationship between abstract non-verbal reasoning and model-based measures in a CCD subsample. Finally, we note that there was no systematic difference in participant accuracy rates, indicating that perceptual decision-making processes remain intact within the CCD cohort, and we removed individuals from analyses who reported a high number of mistakes during confidence judgements or reported to experimenters that they struggled with the task. In the broader literature, metacognition is only weakly correlated with general intelligence and has been reported as contributing independently to academic performance (Ohtani & Hisasaka, 2018). Finally, our neurotypical sample in Experiment 1 and CCD sample in Experiment 2 showed no correlation between abstract non-verbal reasoning and metacognitive ability.

Taken together, we hypothesize that the corpus callosum and complete information are both necessary components specific to metacognitive sensitivity, and thus efficiency. We predict that metacognitive efficiency across domains outside of perception (domain-general metacognitive efficiency) will be impeded by experimental callosal perturbations; phenomenological and clinical profiles of those with CCD suggest that those with CCD show insensitivity to mistakes within social, abstract, and decision-making domains (Maxfield, et al., 2019; Maxfield et al., 2021). We suggest a broader range of metacognitive tasks be provided under experimental callosal perturbations (such as in Lund et al., 2025) to test whether metacognitive efficiency uniformly reduces across perception, memory, and decision making.

We note some limitations with the design of the current study. Our sample sizes were modest, in part due to the rarity of the patient population. Within individual experiments, this may have led to underpowered effects. Despite this, we demonstrate consistent and replicable effects that are robust to both task variation and the method of task delivery. While this improves confidence in our general results, there may have been more subtle metacognitive changes present that we were unable to detect. In addition, not all CCD diagnoses in Experiment 1 and Experiment 3 could be radiologically confirmed; excluding these participants in supplementary analyses had a negligible impact on our results in Experiment 1, but reduced group-differences of the binocular trials in Experiment 3 (Supplementary Materials). Secondly, it is important to note that our aims were to quantify metacognitive differences between groups, and this involved common assumptions about the relationship between first-order performance (*d*’) and metacognitive sensitivity (meta-*d* ’; Maniscalco & Lau, 2012). The assumptions that components of metacognitive efficiency (meta-*d*’/*d*’) are bias-free have been challenged (Guggenmos, 2021), and other frameworks exist that make alternative assumptions about the relationship between performance and sensitivity (Dayan, 2022; De Martino et al., 2013). Relatedly, we note that while we obtained excellent linear and posterior recovery our data & model, the recovered parameters were more conservative with respect to our original estimates. This is due to the hierarchical nature of the fitting process which maximizes sample power by shrinking parameters closer to the mean (Piray et al., 2018). Our smaller samples are more at risk of a shrinkage bias on recovery. Third, staircase procedures can also introduce higher confidence scores in inattentive participants who end up with lower difficulty (Sarna et al., 2025). However, we found no evidence for group differences in our staircasing procedures, improving experimental confidence in the validity of observations within the RDK. Finally, we fit both coherence trials together in the RDK task to extract overall efficiency and sensitivity metrics per group. Fitting coherences together, rather than separately, can lead to the inflation of metacognitive sensitivity parameters (Rahnev & Fleming, 2019). We find that coherence influences overall metacognitive sensitivity, such that low coherence leads to lower sensitivity compared to fitting both together, which in turn has lower sensitivity compared to high coherence trials alone (see Figure S5). Future analysis of metacognitive efficiency in CCD populations should endeavor to use larger samples to allow fine-grained analysis of this computational nuance.

In summary, we find that CCD leads to systematic and enduring deficits in metacognitive efficiency in visual metacognition tasks, shedding light on the importance of the corpus callosum and overall connectivity in supporting metacognition in the neurotypical brain. We hypothesize that domain-general metacognition is a phenomenon that in-part arises from the increased efficiency of informational transfer between hemispheres supported by the corpus callosum.

## Data & Code

Data and analysis code is freely available here: https://github.com/Brain-Development-and-Disorders-Lab/Barnby_etal_2026_ccd_impairs_metacognition

## Acknowledgements

The authors would like to thank the participants for their time in completing these studies and the family support groups Australian Disorders of the Corpus Callosum (AusDoCC), and the National Organization for Disorders of the Corpus Callosum (NODCC) for their help in recruitment and support of our research. The authors also thank Lisa MacKenzie and Tiffany Earle for their efforts in participant enrolment and liaison. This research was supported by the Australian Research Council (DP210101712 to LJR), as well as laboratory startup funds from Washington University in St Louis (LJR), by the Max Planck Society (PD) and the Humboldt Foundation (PD), the FENS-Kavli Network of Excellence (000-001 to JMB), the Wellcome Trust (WT228268/Z/23/Z to JMB) and the Centre for AI and Machine Learning at the University of Western Australia (JMB).

