## Supplementary Materials for "Corpus Callosum Dysgenesis impairs metacognition: evidence from multi-modality and multi-cohort replications"

**Calibration Assessment**

We first ensured that calibration across all experiments were successful. The *k*-value distributions used for the CCD and NT groups in the modified and unmodified RDK tasks are presented In Table S1; in summary, the mean *k*-values of the CCD cohort were no different from those of the NT cohort in all instances (p >> 0.05, two-sided t-test).

**Table S1**: $k_{\text{med}}-$value distributions and between-group statistical test outcomes in the main trials for the participant groups across all experiments.

| **Calibration** | **CCD** | **NT** | **t-value** | **p** |
| --- | --- | --- | --- | --- |
| Experiment 1 | 0.13 ± 0.10 | 0.09 ± 0.06 | 1.32 | 0.22 |
| Experiment 2 | 0.17 ± 0.09 | 0.14 ± 0.13 | 0.49 | 0.63 |
| Experiment 3: Binocular | 0.21 ± 0.03 | 0.22 ± 0.03 | -0.00 | 0.99 |
| Experiment 3: Monocular | 0.21 ± 0.03 | 0.22 ± 0.03 | 0.14 | 0.89 |
| Experiment 3: Lateralized | 0.24 ± 0.03 | 0.22 ± 0.03 | 1.93 | 0.07 |

There was a marginal impact of between-environment effects on calibration for CCD participants (F=3.17, p=0.06) and a significant impact for NT participants (F=25.89, p<0.001). Tukey test post-hoc analysis with family-wise correction showed a significant difference between VR and computer-based environments for CCD participants (difference = 0.09, p=0.047) and NT participants (difference = 0.13, p<0.001). NT participants also showed a difference between VR and MRI-based environments on calibration (difference = 0.07, p=0.02).

Separate calibrations were conducted for each eye during the calibration in the VR environment. There was an overall within-group effect for presentation type on *k*_med_ values for CCD (F=3.85, p=0.04) but not NT participants. Post-hoc contrasts showed that this was driven by a significant difference between Binocular and Lateralized presentation type (t=2.29, p=0.03) and Monocular and Lateralized presentation type (t=2.51, p=0.02) within CCD participants; Lateralized trials led to higher *k*_med_ values, signifying that CCD participants found these trials more difficult versus the other presentation types. There was no significant group-by-presentation-type interaction. For a visualization of the calibration staircase procedures, see Figure S1.

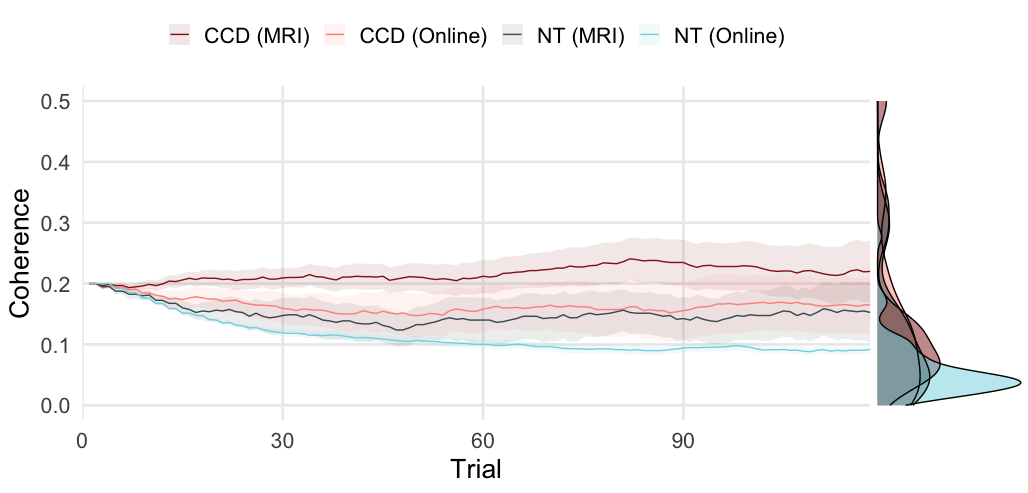

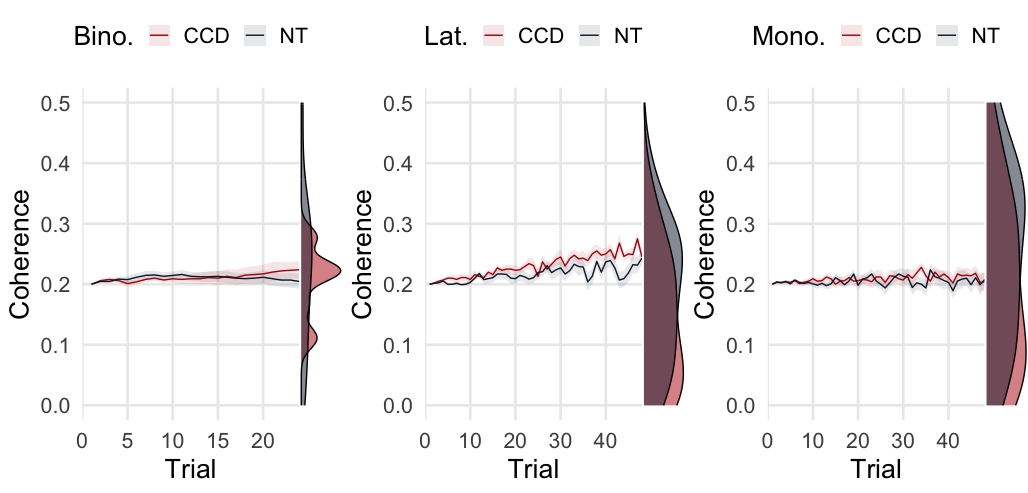

**Figure S1. Calibration outcomes across all experiments.** Graphs depict the group-wise averages of the calculated $k_{\text{med}}$ (coherence) values at each trial during the calibration phase of the experiment using a staircase procedure. Marginal distributions to the right of each graph depict the group distribution of the final coherence value used for the main trials in the RDK tasks. *Top****.*** Experiment 1 and 2 showed no meaningful difference in final $k_{\text{med}}$ values between groups. *Bottom*. No group difference was found in calibration $k_{\text{med}}$ values between groups within conditions.

### **The effect of the task structure and testing environment on behavior**

We performed sensitivity checks to ensure task environment did not influence performance. Initial comparisons of participant accuracy and confidence across the different coherence levels were performed using a two-tailed Welch’s t-test (β = 0.2 and α = 0.05) in R.

A final linear mixed-effects analysis was conducted to ascertain whether the VR modifications and differences in the testing environment (i.e., undertaking the RDK while having an MRI) notably impacted participant accuracy and/or confidence. It was determined that those who participated in the VR modified version of the task had reduced accuracy rates overall (β_VR_ = -0.09, [95% CI = -0.16, -0.02]; p = 0.014), however, there was no difference on participant accuracy across the groups that completed the unmodified version of the task (β_MRI_ = 0.00 [-0.07, 0.08], p = 0.962), indicating that participant performance was consistent across testing environments. Furthermore, the accuracy rates of the CCD participants collectively across the different versions of the tasks were on par with their NT counterparts (β_CCD_ = -0.03, [-0.09, 0.03]; p = 0.350), signifying that the calibration procedure was effective in normalizing the difficulty of the task between the two groups.

This analysis was repeated for participant confidence ratings; confidence ratings were converted into z-scores to improve the comparability of these scores across the modified and unmodified RDK tasks. The task modification was not found to affect participant confidence significantly (β_VR_ = -0.10 [-0.43, 0.22], p = 0.537), nor did the testing environment (β_MRI_ = -0.19 [-0.53, 0.15], p = 0.264); there was no difference in overall confidence ratings between the two groups (β_CCD_ = 0.16 [-0.12, 0.45], p = 0.256). No substantial change was noted in the latter outcomes when the analysis was re-run with the confidence ratings from the modified RDK task excluded.

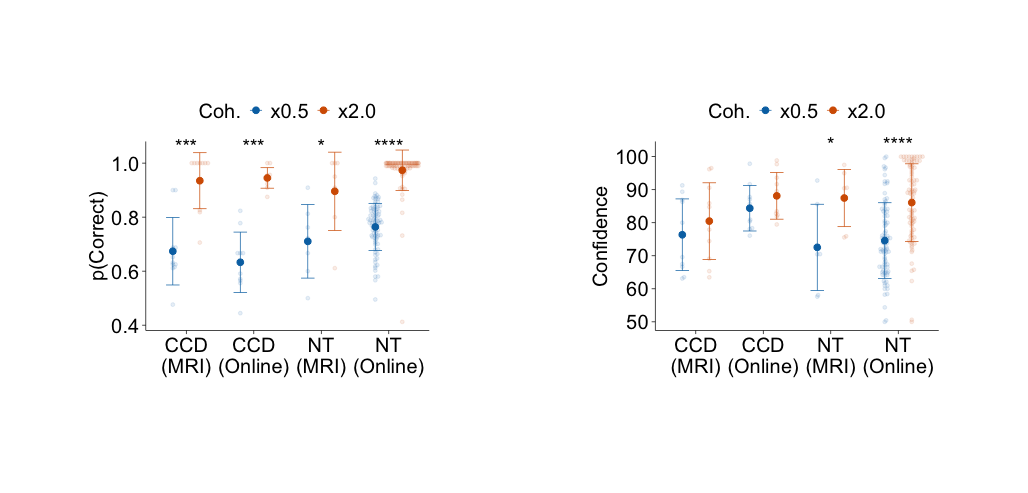

**Figure S2. Raw Behavioral Differences In Experiment 1 and 2.** *Left panel:* All groups showed sensitivity to dot coherence; in all cases participants increased in their accuracy at higher coherence. *Right panel:* Only NT participants calibrated their confidence in line with increased dot coherence. *=p<0.05, **=p<0.01, ***=p<0.001, ****=p<0.0001. Coh. = Dot Coherence ($k_{\text{med}}$). All tests were paired Kruskal-Wallis tests.

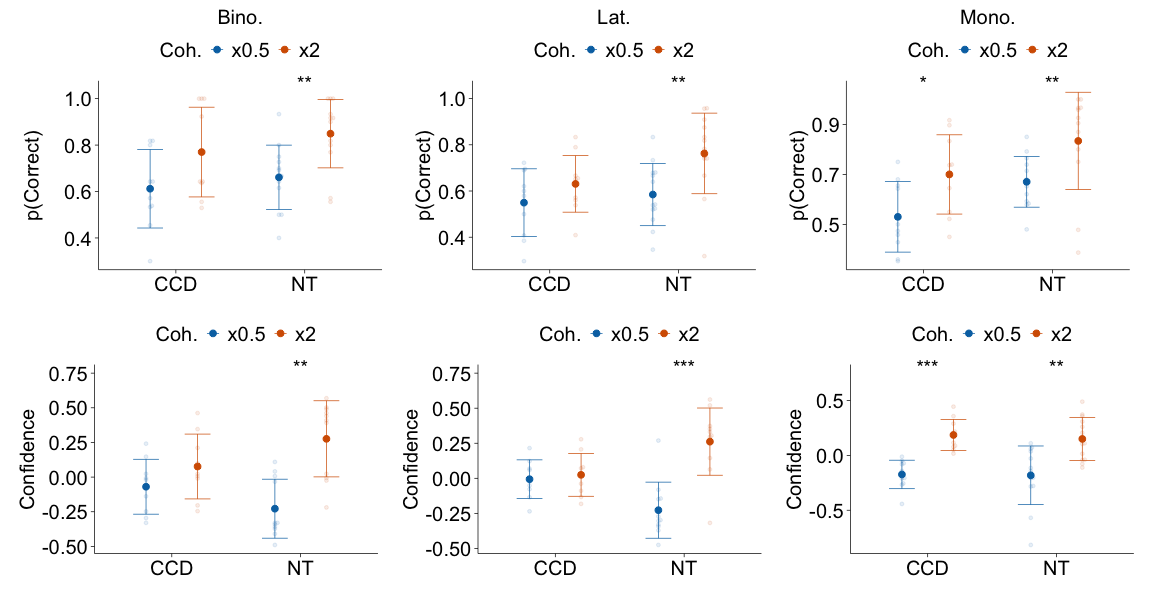

**Figure S3. Raw Behavioral Differences In Experiment 3.** *Left panel:* All groups showed sensitivity to dot coherence; in all cases participants increased in their accuracy at higher coherence. *Right panel:* Only NT participants calibrated their confidence in line with increased dot coherence. *=p<0.05, **=p<0.01, ***=p<0.001, ****=p<0.0001. Coh. = Dot Coherence ($k_{\text{med}}$). All tests were paired Kruskal-Wallis tests.

**Reaction Time Assessment**

We analyzed reaction time as a function of dot coherence, correctness, and group in trials where confidence was not recorded in Experiment 1 and 2 (see Table S2 and S3). This analysis was not possible in Experiment 3 as every response was also tied to a confidence judgement. We address this after trials in Experiment 1 and 2 which did not include a confidence judgement.

We first tested the role of dot coherence and group on log reaction time in each non-confidence trial. We used linear random effect models (random effect of participant ID) to test main effects and interactions. In experiment 1, there was a main effect of group (NT vs CCD; t=-5.56, p<0.001) and coherence (High vs Low; t=-21.77, p<0.001). These terms did not interact. In experiment 2, these findings were replicated; there was a main effect of group (NT vs CCD; t=-2.62, p=0.02) and coherence (High vs Low; t=-3.83, p<0.001). There was no interaction in terms.

We next tested the role of correctness (1 or 0) and group on log reaction time in each non-confidence trial. We used linear random effect models (random effect of participant ID) to test main effects and interactions. In experiment 1, there was a main effect of group (NT vs CCD; t=-6.06, p<0.001) and correctness (1 vs 0; t=-18.26, p<0.001). These terms did not interact. In experiment 2 main effects were replicated; there was a main effect of group (NT vs CCD; t=-3.55, p=0.001) and correctness (1 vs 0; t=-3.00, p=0.003). These terms also interacted (t=3.21, p=0.001), such that those with CCD were relatively faster on correct vs. incorrect trials, suggesting incorrect trials led to slower, more hesitant responses, which were not as prevalent in the NT sample. This may suggest CCD individuals, despite their equivalent accuracy to NTs, required longer under uncertainty to make a decision.

**Table S2. Reaction time (s) and sd split by coherence and group in non-confidence trials.**

| Mean s [sd] | **CCD** | | **NT** | |
| --- | --- | --- | --- | --- |
| **Experiment** | **Low Coherence** | **High Coherence** | **Low Coherence** | **High Coherence** |
| 1 | 1.44 [0.78] | 1.25 [0.55] | 0.68 [0.31] | 0.57 [0.26] |
| 2 | 1.29 [0.43] | 1.16 [0.53] | 0.85 [0.25] | 0.77 [0.17] |

**Table S3. Reaction time (s) and sd split by correctness and group in non-confidence trials.**

| Mean s [sd] | **CCD** | | **NT** | |
| --- | --- | --- | --- | --- |
| **Experiment** | **Correct (1)** | **Incorrect (0)** | **Correct (1)** | **Incorrect (0)** |
| 1 | 1.26 [0.56] | 1.71 [0.96] | 0.60 [0.27] | 0.78 [0.36] |
| 2 | 1.12 [0.37] | 1.54 [0.71] | 0.79 [0.20] | 0.93 [0.26] |

We next analyzed reaction time as a function of dot coherence, correctness, and group during confidence trials (see Table S4 and S5 for raw scores).

We first tested the role of dot coherence and group on log reaction time in each confidence trial. We used linear random effect models (with a random effect of participant ID) to test main effects and interactions. In Experiment 1, there was a main effect of group (NT vs CCD; t=-5.26, p<0.001) and coherence (High vs Low; t=-11.88, p<0.001). These terms did not interact. In Experiment 2, there was no main effect of group (NT vs CCD; t=-1.88, p=0.07) and main effect of coherence (High vs Low; t=-3.92, p<0.001) on reaction time. There was no interaction between terms. In Experiment 3, we divided the models by presentation type. In binocular trials, none of the main or interaction effects were significant. In lateralized trials, there was no effect of group, although there was a main effect of coherence (High vs Low; t-2.50, p=0.01). None of the interaction terms were significant. There was no significant effect in the monocular trials.

We next tested the role of correctness (1 or 0) and group on log reaction time in each confidence trial. We used linear random effect models (with a random effect of participant ID) to test main effects and interactions. In Experiment 1, there was a main effect of group (NT vs CCD; t=-5.80, p<0.01) and correctness (1 vs 0; t=-7.47, p<0.001). These terms interacted (t=3.03, p=0.002), such that those with CCD were relatively faster on correct vs. incorrect trials. In Experiment 2, these findings were replicated; there was a main effect of group (NT vs CCD; t=-3.55, p=0.001) and correctness (1 vs 0; t=-3.00, p=0.003) on reaction time. There was no interaction between terms. In Experiment 3, we divided the models by presentation type. In binocular trials, there was a main effect of correctness (1 vs. 0; t=2.86, p=0.004), but no main effect of group or a significant interaction. In lateralized trials, there was a main effect of correctness (1 vs. 0; t=2.74, p=0.006), but no main effect of group or significant interaction. None of the interaction terms were significant. There was no significant effect in the monocular trials.’

**Table S4. Reaction time (s) and sd split by coherence and group in confidence trials.**

| Mean s [sd] | **CCD** | | **NT** | |
| --- | --- | --- | --- | --- |
| **Experiment** | **Low Coherence** | **High Coherence** | **Low Coherence** | **High Coherence** |
| 1 | 1.52 [0.89] | 1.08 [0.39] | 0.68 [0.30] | 0.58 [0.28] |
| 2 | 1.20 [0.44] | 1.07 [0.57] | 1.03 [0.43] | 0.68 [0.20] |
| 3 (Binocular) | 2.79 [1.53] | 2.68 [1.27] | 2.30 [0.56] | 2.07 [0.40] |
| 3 (Lateralized) | 2.73 [1.20] | 2.59 [0.80] | 2.08 [0.33] | 2.14 [0.47] |
| 3 (Monocular) | 3.11 [1.93] | 2.76 [1.35] | 2.42 [0.81] | 2.16 [0.47] |

**Table S5. Reaction time (s) and sd split by correctness and group in confidence trials.**

| Mean s [sd] | **CCD** | | **NT** | |
| --- | --- | --- | --- | --- |
| **Experiment** | **Correct (1)** | **Incorrect (0)** | **Correct (1)** | **Incorrect (0)** |
| 1 | 1.21 [0.59] | 1.65 [0.77] | 0.61 [0.29] | 0.79 [0.33] |
| 2 | 1.07 [0.50] | 1.35 [0.52] | 0.82 [0.24] | 0.94 [0.34] |
| 3 (Binocular) | 2.69 [1.40] | 2.80 [1.33] | 2.07 [0.40] | 2.56 [0.79] |
| 3 (Lateralized) | 2.80 [1.45] | 3.11 [1.85] | 2.26 [0.71] | 2.46 [0.65] |
| 3 (Monocular) | 2.62 [1.11] | 2.73 [0.93] | 2.10 [0.37] | 2.20 [0.44] |

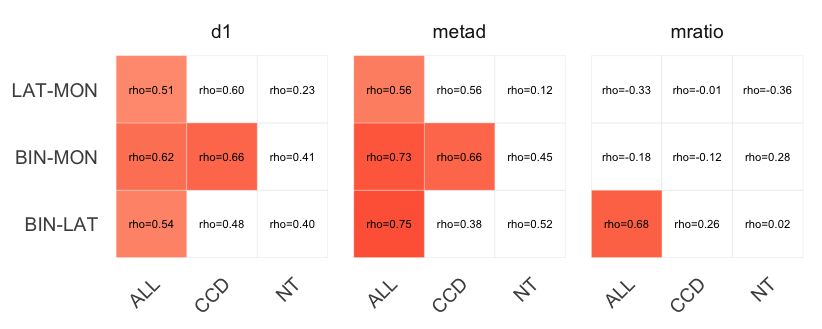

**Figure S4. Within participant Spearman correlations between conditions in Experiment 3.** Correlations (Spearman’s Rho) for each metric are calculated for all participants together (ALL) and by group (CCD, NT). White panels indicate non-significant values (p>0.05). Correlations between m-ratio estimates are a combination of$d'$ and meta-$d'$, and thus their correlation depends on the variability within individuals between conditions. The M-ratio only correlates between binocular and lateralized trials because binocular and lateralized trials show a consistent worsening of d’ across all participants between each type of trial, and thus $d'$ and meta-$d'$ vary together within participants. Given that binocular to monocular and binocular to lateralized trials lead to differing computational impacts between group effects (i.e. $d'$ is the same for NTs but lower for CCDs, and meta-$d'$ is lower for both in binocular to monocular trials), the m-ratios are less likely to correlate.

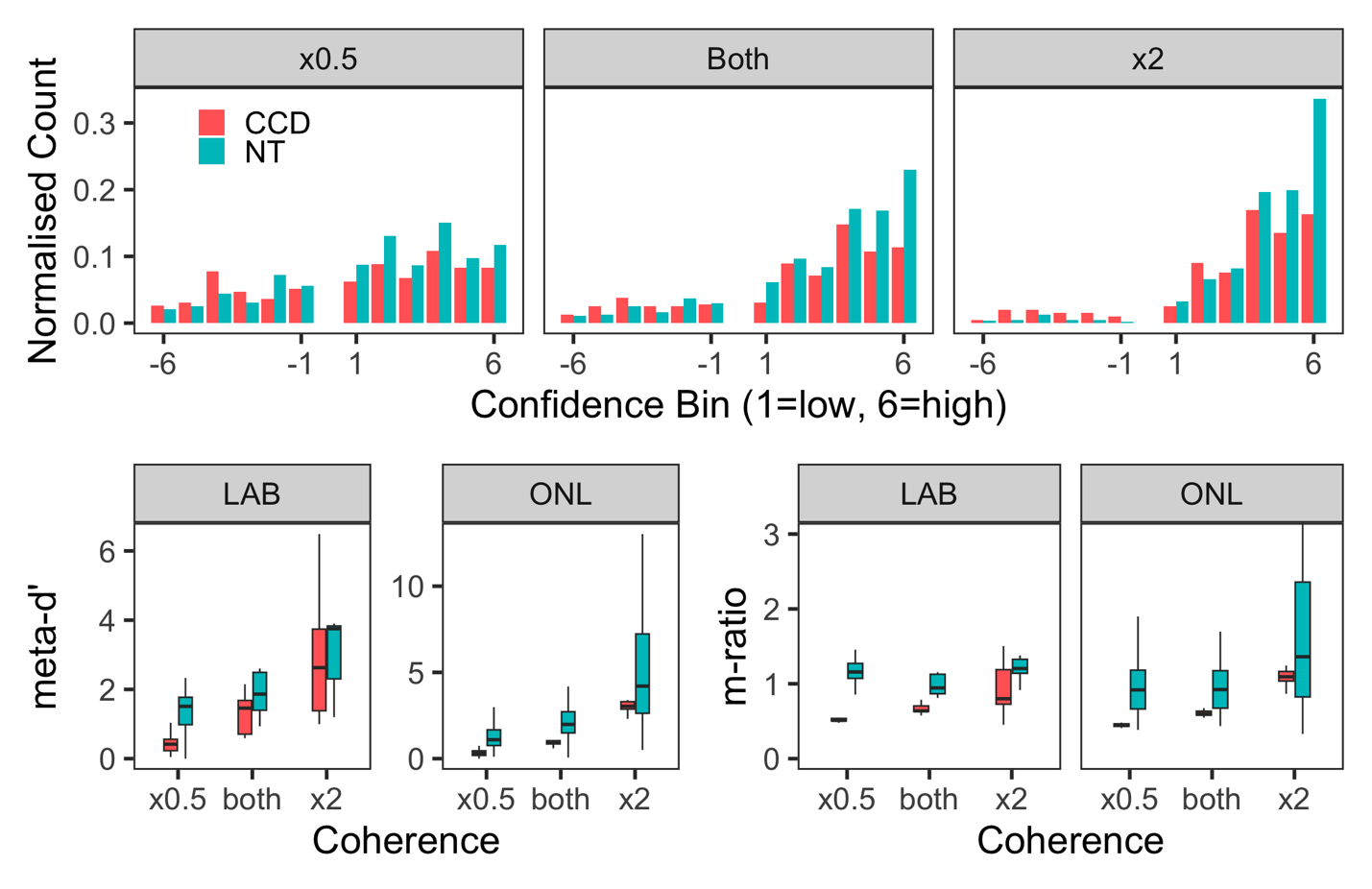
**Figure S5. Spread and distribution of individual confidence attributions, meta-d’, and m-ratio across conditions.** To assess the potential inflation of meta-d’ values that can occur when conditions are mixed within model fitting (Rahnev & Fleming, 2019) we fitted models on data from Experiment 1 and 2 divided by coherence type. In sum, we show that easier trials (coherence = x2) lead to sharper confidence attributions during correct responses, leading to oversampling and inflation of meta-d’ values across the board. The opposite was true for low coherence (x0.5) trials. (Top) The set of three panels shows the normalized count of confidence scores for low (1) and high (6) confidence when an answer was correct (positive values on the x-axis) or incorrect (negative values on the x-axis). We show the spread of confidence attributions when considering all trials together (both), as well as separately depending on coherence (x0.5 or x2). (Bottom) We show the results from model fitting of the individual parameter values for meta-cognitive sensitivity (meta-d’) and metacognitive efficiency (m-ratio) across all trials (both), or when coherences were fitted separately.

**Figure S6. Posterior Predictive Check and Parameter Recovery. (A)** Real (dark blue) and Recovered (gold) confidence counts produced from simulating data within the hierarchical model fitting process used to estimate m-ratio, meta-d and d’ estimates for each experiment and each group. Experiments 1 and 2 used confidence bins that spanned from 50% to 100% in increments of 10% (6 bins in total). Negative values of confidence bins (e.g. -1) indicate confidence assigned to stimuli reported to go left, and positive values of confidence bins the confidence assigned to stimuli reported to go right. **(B)** Pearson r correlations between real and recovered confidence values for each experiment depicted in A. Red line indicates r=0.8. **(C)** Pearson r correlations between real (x-axis) and recovered (y-axis) M-ratio values across all three experiments. Only binocular data are presented for Experiment 3.

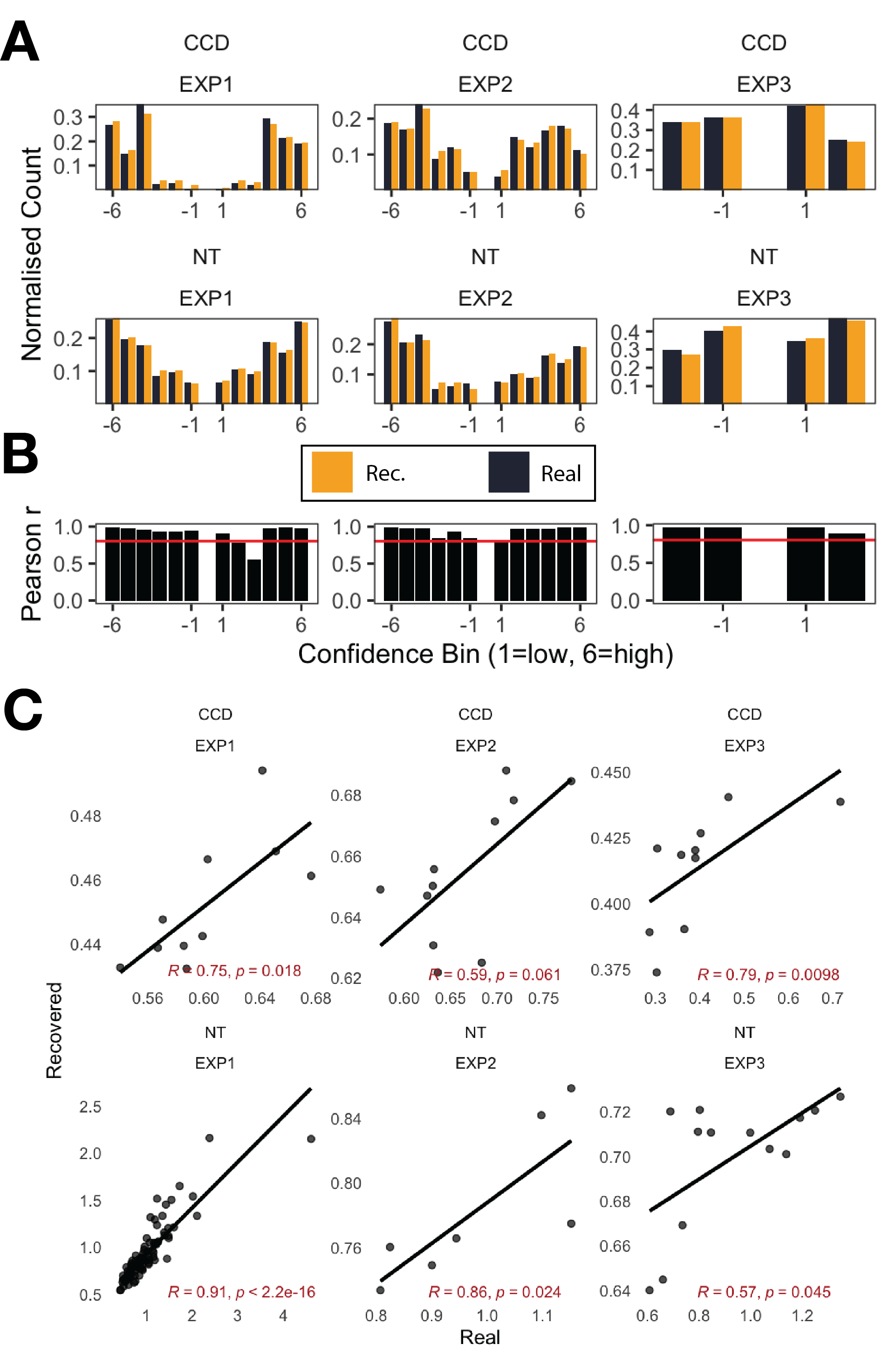

**Task variants and specific modifications**

The first task variant (experiments 1 and 2) utilized identical dimensions and timings described by Rausch & Zehetleitner, 2014 and Bang et al., 2018. Modifications were made to constrain coherent dot motion along the horizontal axis, but the multi-directional distractor dot motion was preserved. The reference direction decision was presented to participants as either the left or right side of the dot aperture. These changes were made in anticipation of task difficulty for CCD participants, and we used a staircase to calibrate dot coherence to control task difficulty in line with previous work (Rausch & Zehetleitner, 2014). When using a 4-button MR-safe handheld controller, experiment 2 pilot testing feedback resulted in simplification of confidence input. Input mappings were re-assigned to match the directionality of the stimulus and horizontal positioning of controller buttons, where the left-most controller button was assigned position 0, and the right-most controller button was assigned position 3 on the controller. Left-ward inputs (such as selecting the left reference direction or decreasing confidence) were assigned the left-most button (index 0), and right-ward inputs (such as selecting the right reference direction or increasing confidence) were assigned the right-most button (position 3). Recording a mistake in the reference selection was assigned to the button at position 1, and submitting a confidence input was assigned to the button at position 2. Source code for the first task variant is available on GitHub: https://github.com/Brain-Development-and-Disorders-Lab/task_rdk/tree/v1.8.0

The second task variant (experiment 3) utilized a rectangular aperture of identical area, with the long edge along the vertical axis. This change was in response to constraints imposed by the VR environment’s field of view. Furthermore, coherent dot motion was constrained along the vertical axis, but the multi-directional distractor dot motion was preserved. Similarly, the reference direction decision was represented by a pair of buttons above and below the aperture. Participants were required to hold the VR controller trigger for 1000ms to make their selection, and they were shown a progress bar overlaid on the selected button during this time. Force feedback was issued through the controller upon selection. Dot motion was presented for 180ms to ensure the integrity of lateralized presentation, and eye-tracking functionality was utilized to enforce central fixation during dot motion presentation. Source code for the second task variant is available on GitHub: https://github.com/Brain-Development-and-Disorders-Lab/task_vr_rdk/tree/v1.3.2

**Exclusion Analysis**

We examined key behavioral and modelling outcomes while excluding participants in Experiment 1 (n=1) and Experiment 3 (n=4) who self-reported, rather than confirmed, their radiological diagnosis.

**Experiment 1**

The task showed good discriminability, such that higher moving dot coherence led to improved accuracy scores in both groups (CCD: t=7.84, es=0.77, p<0.001; NT: t=15.22, es=0.34, p<0.001). There was a group by coherence interaction, such that NT participants scored more highly than CCD participants in low but not high coherence trials (t=3.03, es=0.30, p=0.002).

Both CCD and NT participants increased their mean confidence in line with increasing coherence (CCD: t=4.12, es=0.38, p<0.001; NT: t=27.69, es=0.49, p<0.001). There was a group by coherence interaction, such that NT participants had higher confidence in low coherence conditions versus CCD participants (t=5.77, es=0.46, p<0.001).

Computational modelling revealed that those with CCD were far less efficient in their calibration of confidence relative to their perceptual accuracy ($\mu_{\text{ratio}}^{\text{CCD}}$ = 0.63 vs $\mu_{\text{ratio}}^{\text{NT}}$ = 0.88). There was good evidence that the difference in distributions was outside the null ($\Delta\mu_{\text{ratio}}$=0.30, $U$=0.93, 95%CI[$U$]: 0.90, 0.95).

**Experiment 3**

The VR-based task showed good discriminability in binocular trials, such that higher dot coherence led to improved accuracy scores in both groups (CCD: t=2.32, es=0.27, p=0.02; NT: t=3.90, es=0.30, p<0.001). Analysis of lateralized trials showed the same effect for both groups (CCD: t=2.14, p=0.033; NT: t=4.89, es=0.27, p<0.001). Analysis of monocular trials showed accuracy improved when coherence increased in both CCD and NT individuals (CCD: t=3.72, es=0.30, p<0.001; NT: t=4.91, es=0.27, p<0.001). Analysis of all data together, irrespective of presentation, using random-effect models type found main effects for coherence (t=5.35, es = 0.27, p<0.001) and group (t=2.78, es=0.21, p=0.01) on accuracy.

In binocular trials, both NT and CCD changed their mean confidence in line with higher dot coherence (CCD: t=2.33, es=0.30, p=0.02; NT: t=4.25, es=0.36, p<0.001). Only NT individuals changed their confidence in line with coherence in lateralized trials (CCD: t=1.10, p=0.27; NT: t=5.79, es=0.35, p<0.001). Both CCD and NT participants adjusted their confidence in line with coherence during monocular trials (CCD: t=3.46, es=0.28, p<0.001; NT: t=4.09, es=0.23, p<0.001). Analysis of all data together, irrespective of presentation, using random-effect models type found main effects for coherence (t=3.91, es = 0.21, p=0.001), but not group x coherence interaction.

Computational modelling revealed that those with CCD were less metacognitively efficient ($\mu_{\text{ratio}}^{\text{CCD}}$ = 0.52 vs $\mu_{\text{ratio}}^{\text{NT}}$ = 0.78). There was weak evidence that the difference in distributions was outside the null when excluding participants ($\Delta\mu_{\text{ratio}}$=0.26, $U$=0.71, 95%CI[$U$]: 0.66, 0.74).

**Table S6. Phenotypic subtypes within CCD groups for each experiment.** We note only participants who were included within analyses. To note: 1 participant in Experiment 1 had an unknown phenotype. We excluded them from our analysis to test robustness of our results (Supplementary Materials: Exclusion Analysis).

|  | **Experiment 1** | **Experiment 2** | **Experiment 3** | **Total** |
| --- | --- | --- | --- | --- |
| **CCD** | 3 | 6 | 4 | **13** |
| **ACC** | 6 | 4 | 6 | **16** |
| **Hypoplasia** |  | 1 |  | **1** |
